## Supplemental Materials for "Contribution of Retrotransposition to Developmental Disorders"

### Supplemental Figure 1

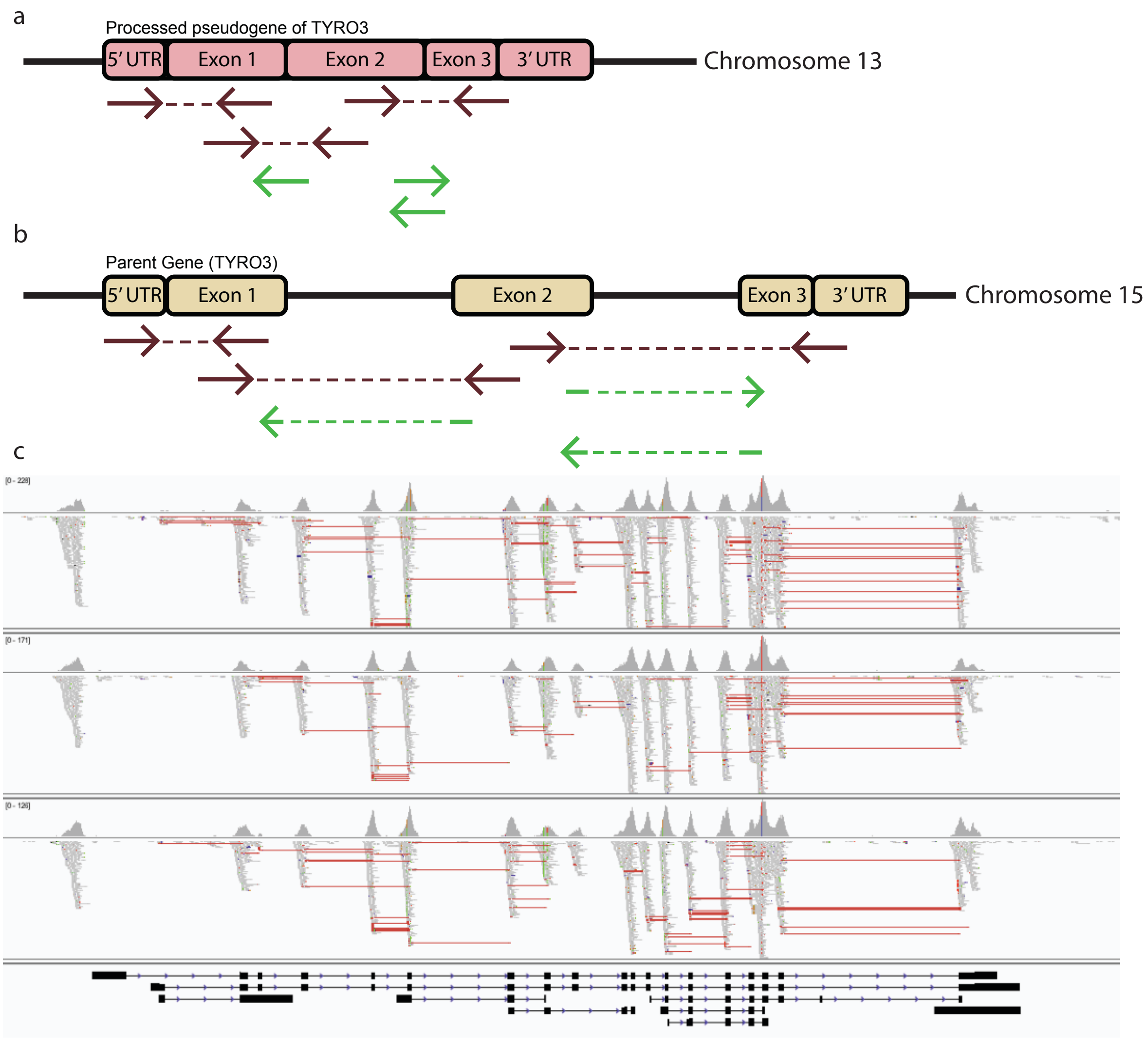


**Identification of processed pseudogenes (PPGs).** Included is an example of the read-pair evidence utilized by our PPG discovery pipeline. Shown throughout the figure is how read evidence supporting the duplication of *TYRO3* aligns to: (**a**) a cartoon example of the PPG, (**b**) a cartoon of the donor gene, and (**c**) a screenshot of read-pair evidence supporting the duplication aligned to *TYRO3* for one trio viewed in the Integrative Genomics Viewer (IGV)[1]. For the cartoons in (**a**) and (**b**), split read pairs (SRPs) are shown in green, with discordant read pairs (DRPs) shown in dark red. When aligned to the duplication, SRPs and DRPs align without gaps and/or clipped sequence. When aligned to the source gene as in (**b**), these reads align further apart than expected or have clipped ends (long dashed lines). This is shown in real data in (**c**), where the red lines represent DRPs that have a larger insert size than the WES library preparation.

**Supplemental Figure 2**

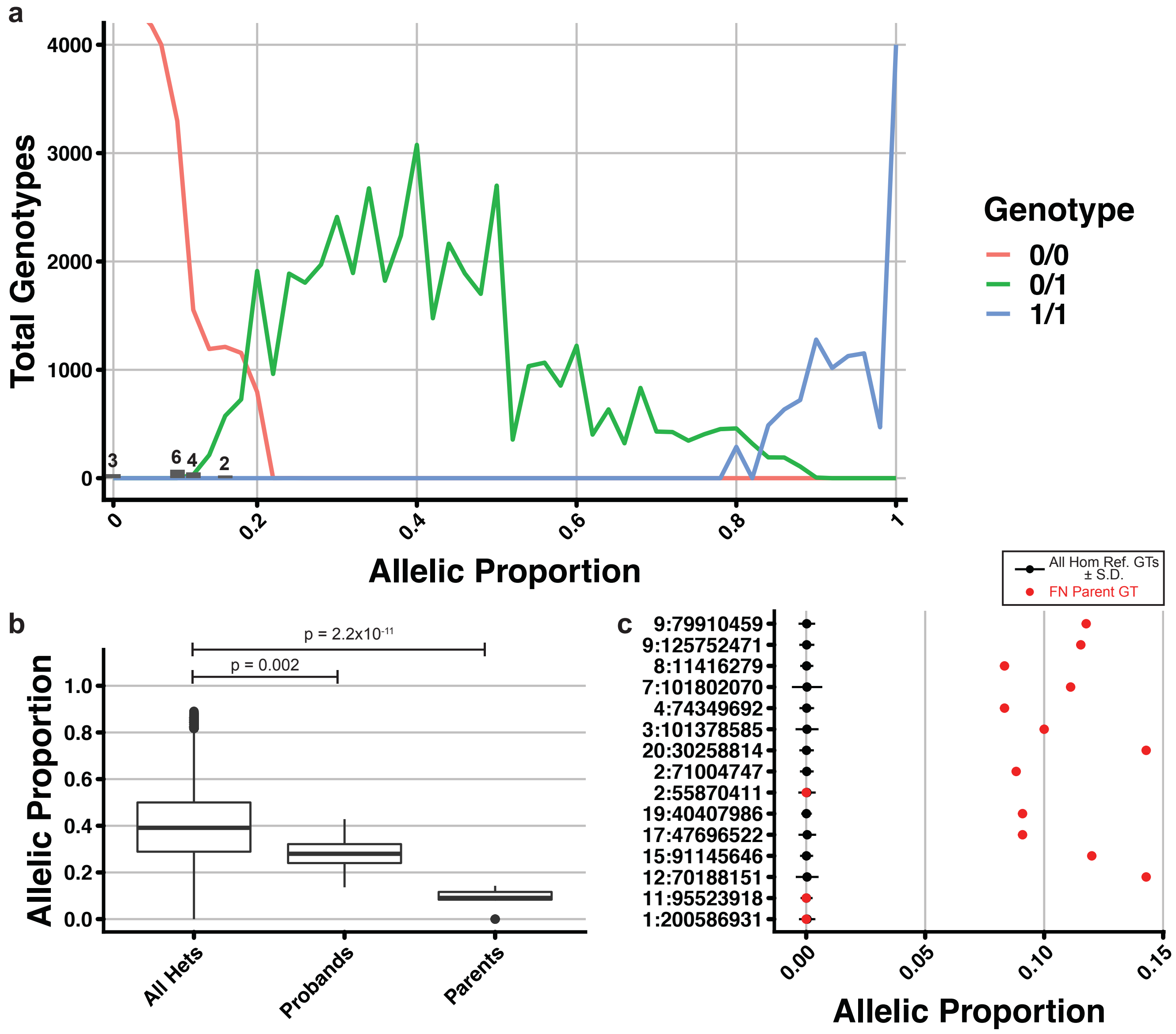


**Allelic proportion of *Alu* variants in the DDD study.** (**a**) Calculation of allelic proportion from a subset of DDD trios. Displayed are a total of 2,532,531 non-fail (./.) genotypes from the 917 *Alu* sites identified in this study with a depth of at least 10X coverage separated into 0.02 allelic proportion bins for 1,000 randomly selected and 15 false negative trios. Allelic proportion was calculated as number of insert supporting reads / total reads spanning the insertion site. Lines are coloured based on the MELT-assigned genotype (hom ref [0/0], het [0/1], hom alt [1/1] as red, green, blue, respectively). Note that, due to the excess of homozygous reference genotypes, the line indicating total homozygous reference sites continues above the y-axis. *Alu* sites with a false negative genotype in a parent (i.e. initially thought to be *de novo*) are shown as grey bars with total number of sites in each bin shown above. As is shown by this plot, false negative genotypes do fall on the extreme end of true heterozygous genotypes, but also fall within other sites which are deemed homozygous reference by MELT genotyping. (**b**) Comparison of allelic proportion. Shown are boxplots of aggregate allelic proportion for all heterozygous variants re-genotyped in (**a**; All Hets), 15 heterozygous variants called as *de novo* but on manual inspection were likely inherited (Probands), and corresponding parents to this subset of probands (Parents). P-values above boxplots were calculated via a Wilcoxon rank-sum test. While both results are statistically significant and our potentially mosaic loci differ in allelic proportion to others in the genome, the true difference from the median allelic proportion of all heterozygous variants for parental genotypes (difference = 0.30) is clearly larger than that for MELT-called heterozygous probands (difference = 0.11). (**c**) Assessment of genotyping error at false negative loci. Black points (lines ± S.D.) are mean allelic proportion for all homozygous reference genotypes at each of the 15 assessed false negative *de novo* loci. Red points indicate the allelic proportion in each of the 15 false negative parents at the locus indicated on the y-axis. In this case, 12/15 (80%) false negative parents showed a difference from the homozygous allelic proportion distribution (z-test $p < 0.05$). Additionally, the individuals with homozygous reference genotypes at each locus do not show higher error rates and are closely centered around the expected proportion of 0.

**Supplemental Figure 3**

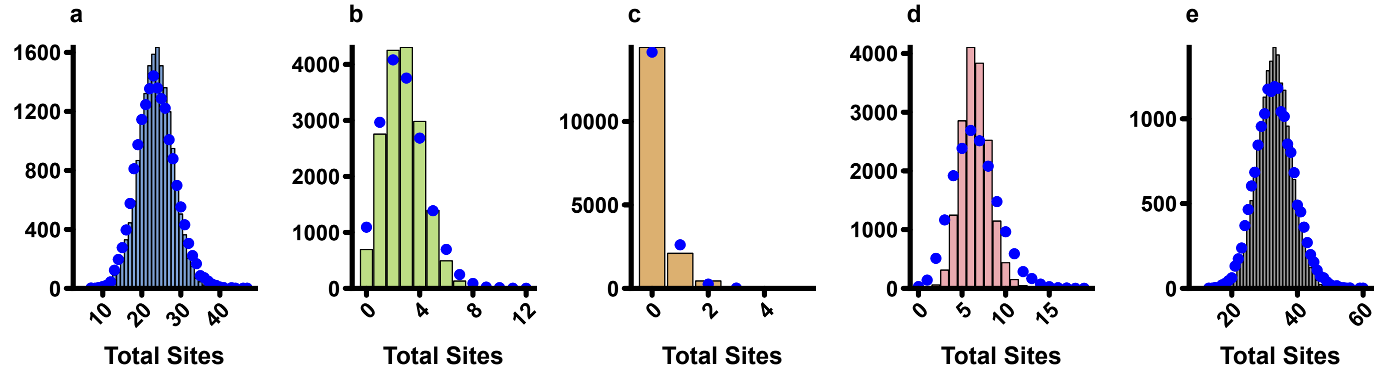


**Poisson distribution projected onto allele counts per individual.** Shown are identical plots to main text Fig. 1a-e for *Alu* (a), L1 (b), SVA (c), PPGs (d), and all RT events (e), but with Poisson distributions based on the mean number of sites per individual projected onto each (shown as blue points).

**Supplemental Figure 4**

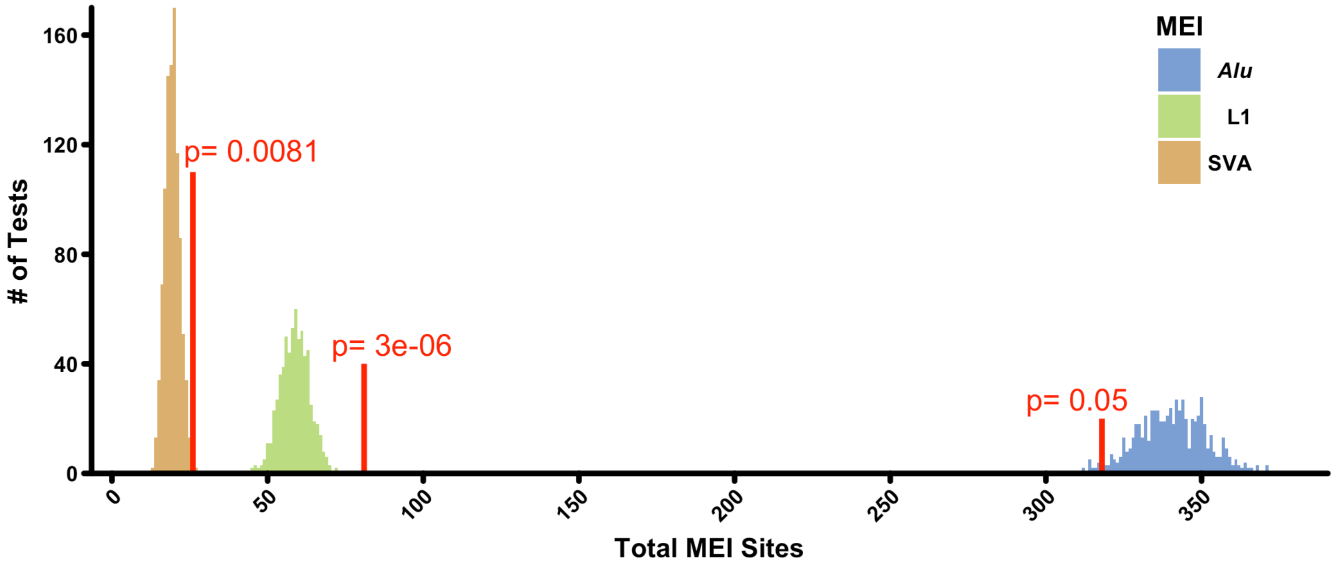


**Downsample of DDD to 1KGP size.** Shown are histograms for DDD data randomly down-sampled 1,000 times to the population size of the 1KGP cohort for all three MEI types (*Alu* – blue, L1 – green, and SVA – orange). Red lines indicate the total number of sites in the 1KGP within WES bait regions, with p-values based on z-score distributions for each MEI class shown above.

**Supplemental Figure 5**

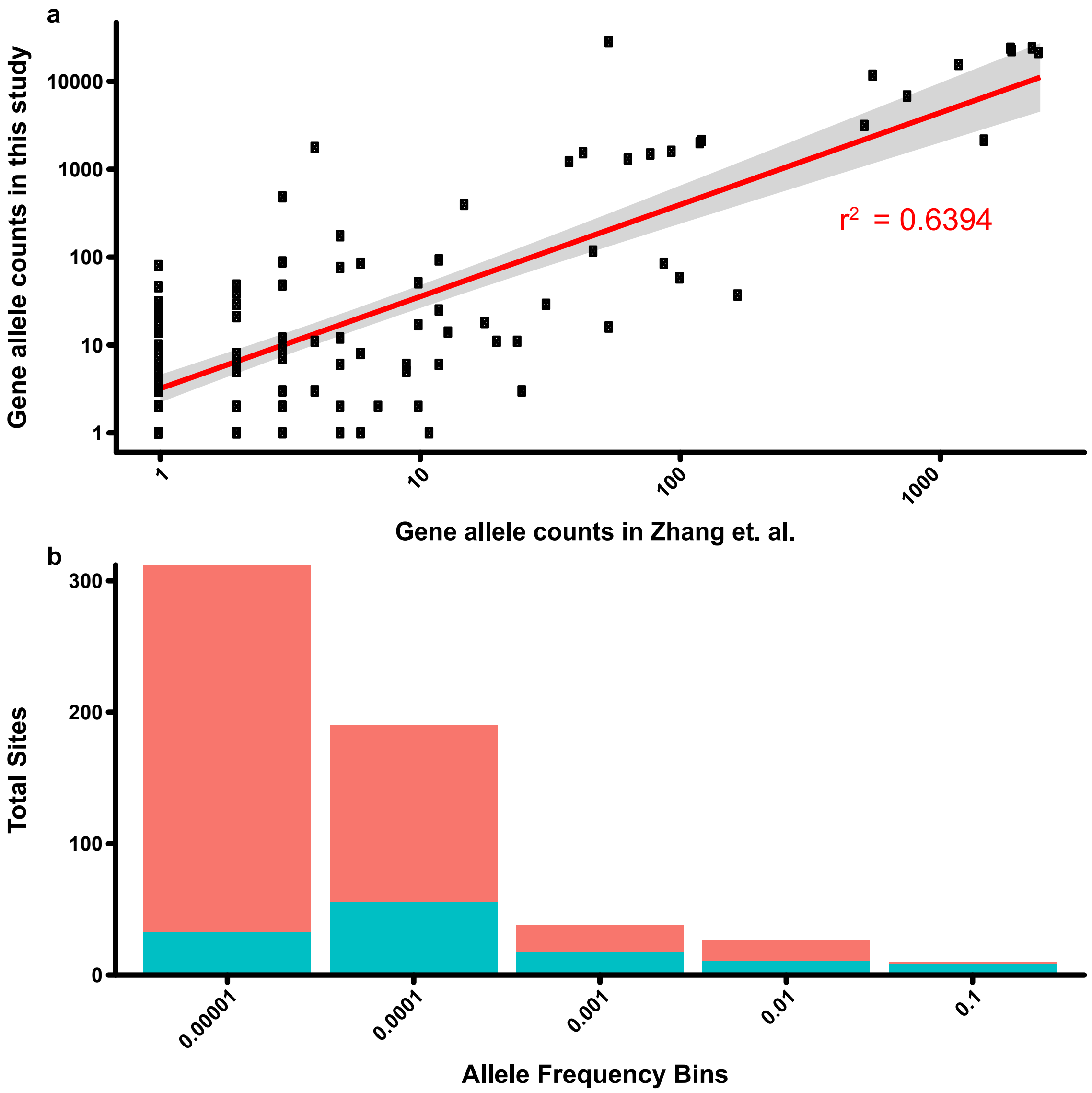


**Comparison of Zhang et. al.[2] and this study.** (**a**) Allele count comparison between Gardner & Zhang. Points represent total number of individuals in which a PPG was observed for Zhang et. al.[2] (x-axis) and this study (y-axis). Regression line is given in red, with calculated $r^2$ listed below. (**b**) Counts of known (light blue) and unknown (pink) PPGs separated into $\log_{10}$ allele frequency bins. The majority of rare PPGs (AF < 0.001) were identified only in this study.

### Supplemental Figure 6

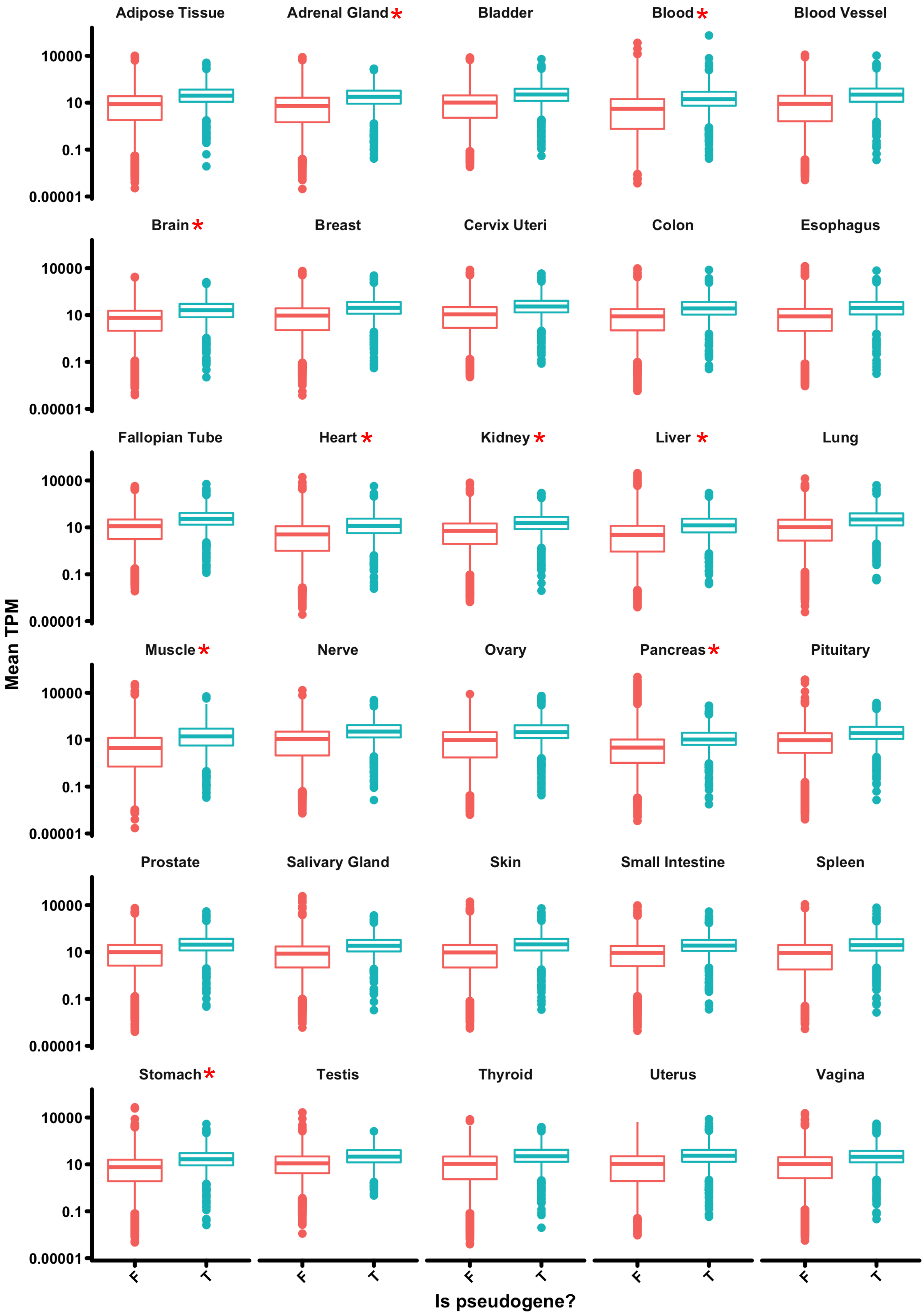


**Donor gene expression in GTEx[3].** Shown for each tissue type are violin plots of the expression in TPM of donor genes that gave rise to duplications (T – light blue) and did not give rise to duplications (F – light red). Within each tissue, genes which gave rise to duplications have TPM values which are significantly higher (Wilcoxon rank-sum p-value < $1x10^{-3}$) than those of their non-duplicated counterparts. * - indicates tissues which have a Wilcoxon rank-sum p-value < $1x10^{-3}$ when compared to testis and ovary donor gene expression.

**Supplemental Figure 7**

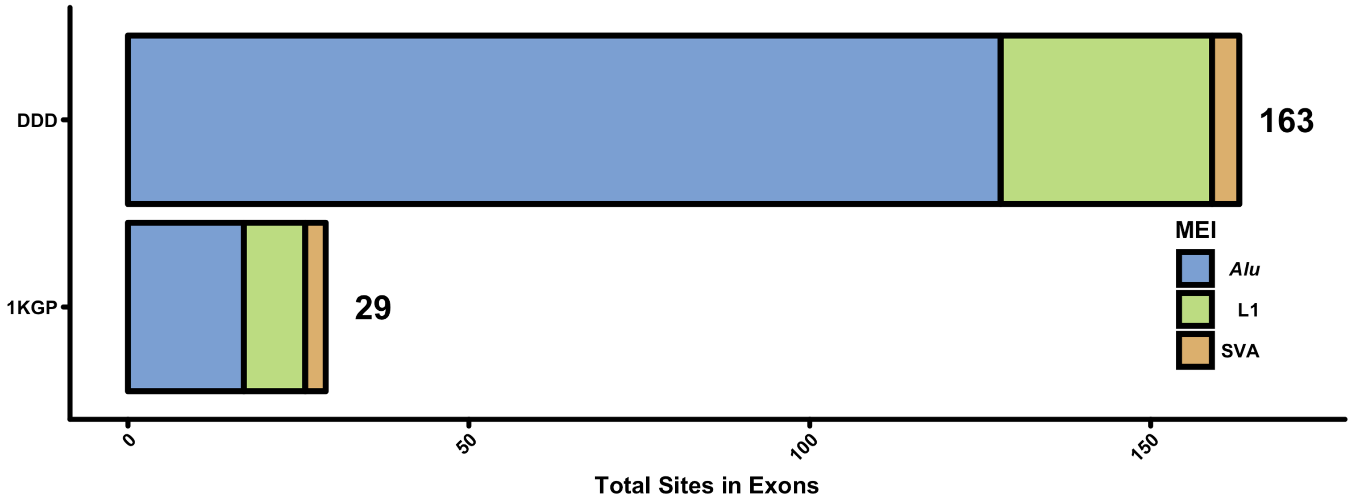


**Total exonic variants identified in 1KGP and DDD.** Shown are total exonic variants as identified in the 1KGP compared to those identified in this study. Each bar plot is divided into the three ME classes analysed as part of this study (*Alu*, L1, and SVA).

**Supplemental Figure 8**

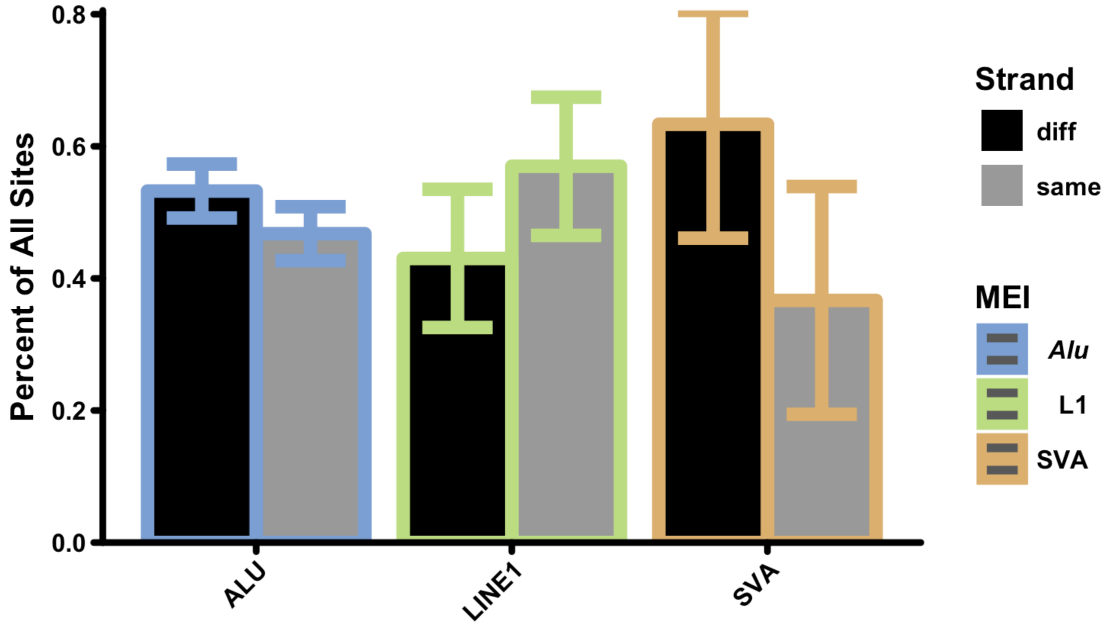


**Strand bias data for intronic MEI sites.** Percentage of all MEIs that are either in the same orientation (same – grey) or in opposite orientation (diff – black) as their host gene for all three ME types (*Alu* – blue, L1 – green, and SVA – orange). None of the differences shown are statistically significant at p = 0.05.

**Supplemental Figure 9**

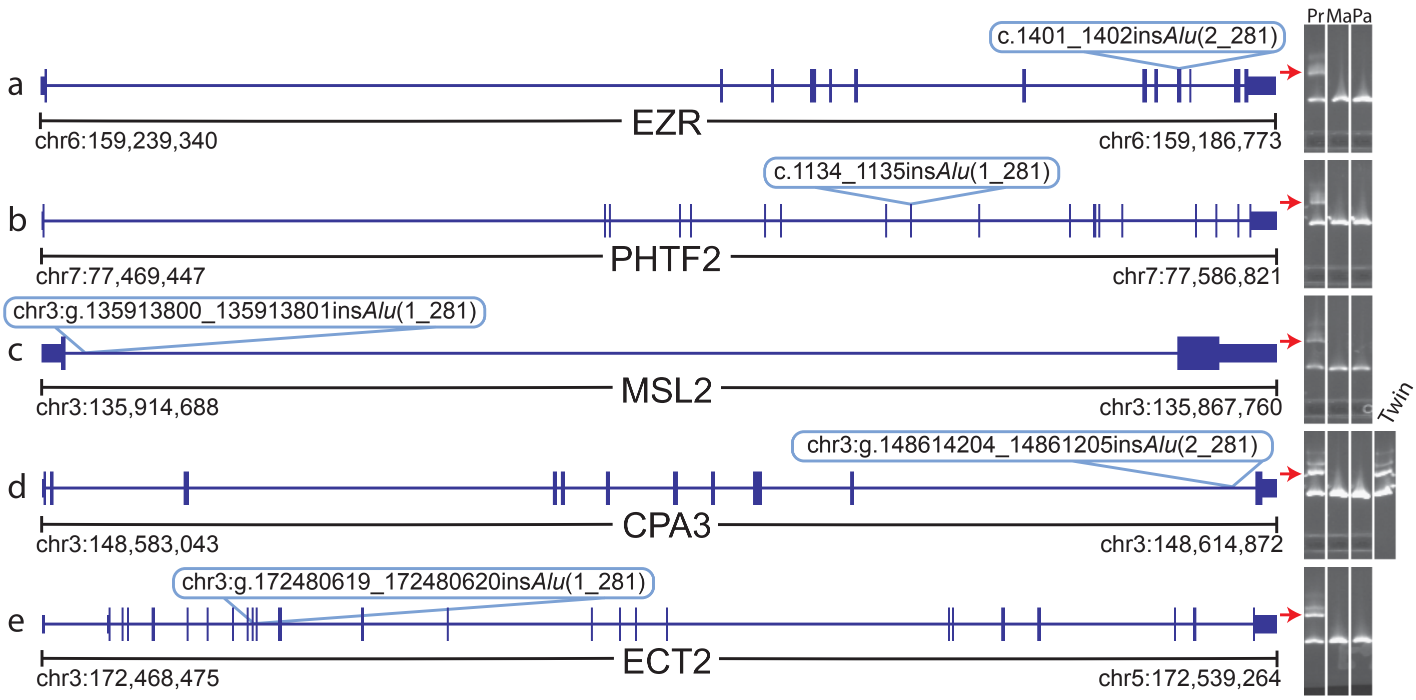


**Non-clinically relevant de novo MEIs.** Shown are plots identical to those in main text Fig. 3a-d, except for *de novo* MEIs that are not clinically actionable. Each gene is shown with site of impact with the colour of the bubble indicating the causal RT type. For CPA3 (**d**), additionally shown is the *de novo* MEI found in the proband's twin to the right of the proband (Pr), mother (Ma), and father (Pa).

**Supplemental Figure 10**

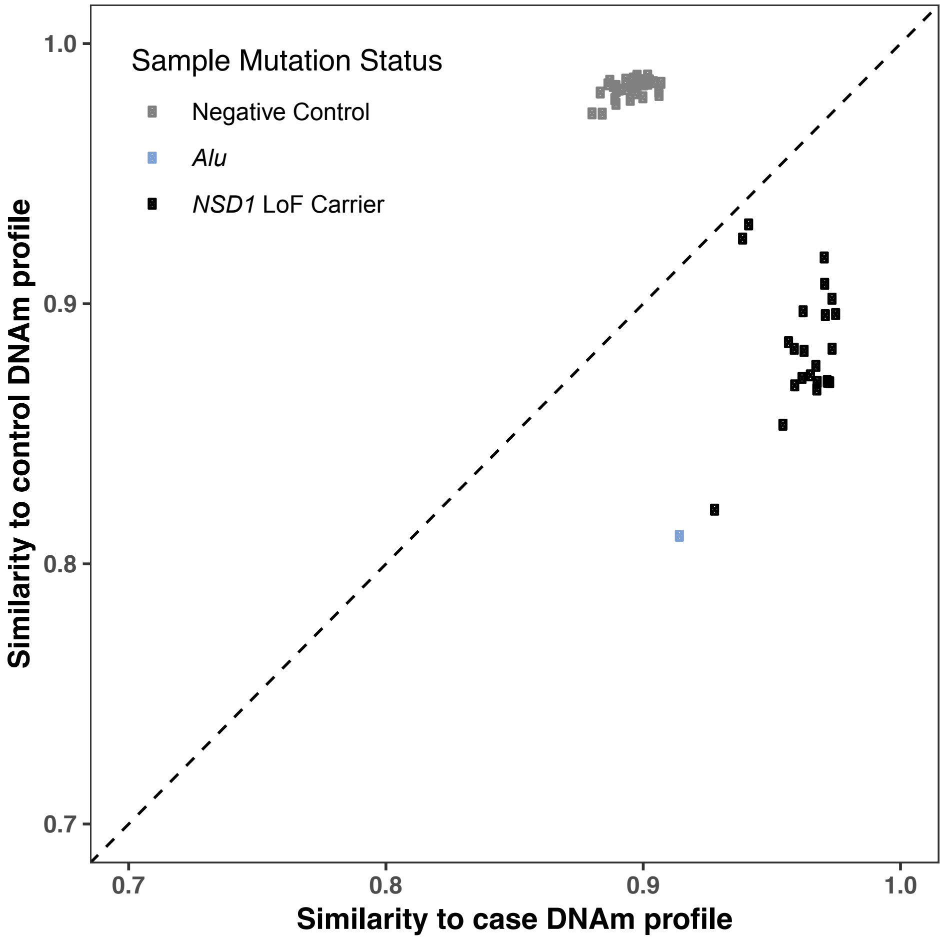


**Methylation analysis of individual with NSD1 loss of function *Alu* insertion.** Shown are Pearson correlation scores (see methods) to the methylation profile of NSD1 LoF mutation carriers (y-axis) and control individuals (x-axis). The individual with the likely-causative NSD1 *Alu* variant identified in this study is shown in light blue, while case and control individuals from Choufani, et al. [4] are shown as black and grey points, respectively.

**Supplemental Figure 11**

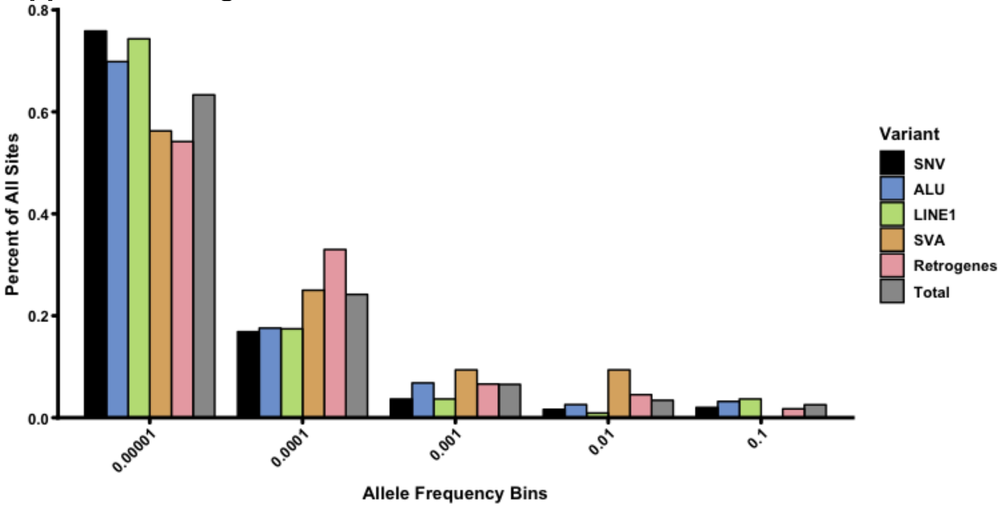


**Allele frequency plot in >10X coverage bait regions.** Allele frequency plot for all four RT classes, total, and SNVs for comparison. Identical to the plot in main text Fig. 1F, but only for variants within the DDD accessible genome mask (see methods).

**Supplemental Table 1**

| ***Alu*** | | | | | |
|---|---|---|---|---|---|
| | | WGS | | | |
| | | 0/0 | 0/1v | 1/1 | ./. |
| WES | N.P. | 234559 | 65405 | 16006 | 221 |
| | N.D. | 760 | 65 | 47 | 7 |
| | 0/0 | 2295 | 123 | 4 | 1 |
| | 0/1 | 12 | 944 | 61 | 1 |
| | 1/1 | 0 | 26 | 258 | 0 |
| | ./. | 51 | 3 | 1 | 0 |

| **L1** | | | | | |
|---|---|---|---|---|---|
| | | WGS | | | |
| | | 0/0 | 0/1 | 1/1 | ./. |
| WES | N.P. | 36944 | 7232 | 2236 | 44 |
| | N.D. | 158 | 9 | 0 | 0 |
| | 0/0 | 458 | 8 | 0 | 0 |
| | 0/1 | 1 | 132 | 8 | 0 |
| | 1/1 | 0 | 0 | 20 | 0 |
| | ./. | 0 | 0 | 0 | 0 |

| **SVA** | | | | | |
|---|---|---|---|---|---|
| | | WGS | | | |
| | | 0/0 | 0/1 | 1/1 | ./. |
| WES | N.P. | 12758 | 2624 | 137 | 25 |
| | N.D. | 411 | 7 | 1 | 0 |
| | 0/0 | 55 | 1 | 0 | 0 |
| | 0/1 | 0 | 1 | 0 | 0 |
| | 1/1 | 0 | 0 | 0 | 0 |
| | ./. | 0 | 0 | 0 | 0 |

**Genotype matricies of matched WGS and WES data.** Shown individually for each ME are genotype matrices for sites identified in both WES (left) and WGS (top) data. N.P. (Not possible) are genotypes that were <10x coverage in the matched WES sample and thus were not expected to be identifiable by MELT. N.D. (Not detected) are genotypes that were identified in the WGS data that were >10x coverage in the matched WES sample and should have been identified by MELT. Highlighted in yellow are the matched genotypes used for the genotype accuracy calculation listed in the main text.

**Supplemental Table 3**

| Insertion Coord. | RT Type | HGNC Gene ID | HPO Terms | HPO Phenotypes |
|---|---|---|---|---|
| **chr3:9495459** | ***Alu*** | **SETD5** | **HP:0000220\|HP:0000431\|HP:0000455\|HP:0000846\|HP:0001161\|HP:0001263\|HP:0001830\|HP:0002205\|HP:0002342\|HP:0006136\|HP:0008915\|HP:0100000** | **Adrenal insufficiency\|Bilateral postaxial polydactyly\|Broad nasal tip\|Childhood-onset truncal obesity\|Early onset of sexual maturation\|Global developmental delay\|Hand polydactyly\|Intellectual disability\|moderate\|Postaxial foot polydactyly\|Recurrent respiratory infections\|Velopharyngeal insufficiency\|Wide nasal bridge** |
| **chr5:176638159** | ***Alu*** | **NSD1** | **HP:0000238\|HP:0000256\|HP:0001249\|HP:0001250\|HP:0001252\|HP:0001382\|HP:0000729\|HP:0010864\|HP:0000540\|HP:0000733\|HP:0001763\|HP:0200006\|HP:0001263** | **Hydrocephalus\|Intellectual disability\|Joint hypermobility\|Macrocephaly\|Muscular hypotonia\|Scoliosis\|Autistic Behaviour\|Severe ID\|Hypermetropia\|Steryotypy\|Pes planus\|Slanting of the palpebral fissures\|Global developmental delay** |
| **chr12:46246325** | **L1** | **ARID2** | **HP:0000154\|HP:0000176\|HP:0000316\|HP:0000347\|HP:0000475\|HP:0000520\|HP:0000586\|HP:0000960\|HP:0001476\|HP:0001601\|HP:0002020\|HP:0002209\|HP:0008434\|HP:0008872** | **Broad neck\|Delayed closure of the anterior fontanelle\|Feeding difficulties in infancy\|Gastroesophageal reflux\|Hypertelorism\|Hypoplastic cervical vertebrae\|Laryngomalacia\|Micrognathia\|Proptosis\|Sacral dimple\|Shallow orbits\|Sparse scalp hair\|Submucous cleft hard palate\|Wide mouth** |
| **chr5:88100580** | **L1** | **MEF2C** | **HP:0000750\|HP:0000915\|HP:0010864** | **Delayed speech and language development\|Intellectual disability, severe\|Pectus excavatum of inferior sternum** |

**Patient phenotypes and HPO terms.** Listed are HPO terms and matched phenotypes as determined by the referring clinician for patients with an identified, clinically actionable MEI. Insertion coord. Is the Hg19 insertion site as listed in main text Table 2.

**Supplemental Table 4**

| Dataset | MEI | Mask | Mask Size | # of haplotypes | # Seg Sites | Theta (ϴ) | Effective Populaton Size (Ne) | Genome Mutation Rate (u = ϴ / 4*Ne) | Mutation Rate |
|---|---|---|---|---|---|---|---|---|---|
| DDD | *Alu* | Exome | 74.2E+6bp | 34046 | 660 | 59.931 | 10000 | 0.001498274 | 1.00941E-11 |
| DDD | L1 | Exome | 74.2E+6bp | 34046 | 109 | 9.8977 | 10000 | 0.000247442 | 1.66706E-12 |
| DDD | SVA | Exome | 74.2E+6bp | 34046 | 32 | 2.9057 | 10000 | 7.26436E-05 | 4.89412E-13 |
| 1KGP | *Alu* | Noncoding | 11.1E+8bp | 4906 | 8554 | 942.56 | 10000 | 0.023563887 | 1.05854E-11 |
| 1KGP | L1 | Noncoding | 9.6E+8bp | 4906 | 2047 | 225.56 | 10000 | 0.005638915 | 2.93737E-12 |
| 1KGP | SVA | Noncoding | 9.6E+8bp | 4906 | 329 | 36.252 | 10000 | 0.000906303 | 4.72104E-13 |

**MEI Mutation rate calculations.** Raw data for MEI mutation rate estimates with individuals values used as input to the Watterson estimator[5]. Dataset indicates which dataset was used for the calculation (DDD – this study; 1KGP – Gardner et. al[6]) with Mask indicative of the genome mask used to filter sites (see online methods).

**Supplemental Table 5**

| Gene | Patient Decipher ID | Gene Median RPKM | Gene RPKM Percentile |
| --- | --- | --- | --- |
| *SETD5* | 280818 | 10.00 | 65.3 |
| *NSD1* | 259118 | 9.26 | 63.2 |
| *ARID2* | 264759 | 4.36 | 38.5 |
| *MEF2C* | 285645 | 84.37 | 96.7 |

**Fetal Brain Expression Data.** Gene RPKM and percentile values as calculated from fetal brain samples 15 to 37 weeks post-conception from the BrainSpan consortium[7].

**Supplemental Data 1**

**Processed pseudogene (PPG) information sheets.** For each *de novo* PPG identified as part of this study, we have collated: primers used (Supplemental Table 3) sequence of the capillary trace (Read Sequence), the resulting trace aligned to the Hg19 human reference genome (Aligned to Hg19 sequence), a graphical representation of the alignment in the UCSC genome browser[8] and, where PCR amplified both PPG and host gene, the intensity data indicating presence of multiple bands in the capillary sequencing.

### SLC35F2

PCR primers: 2F and 3R
capillary primer: 2F

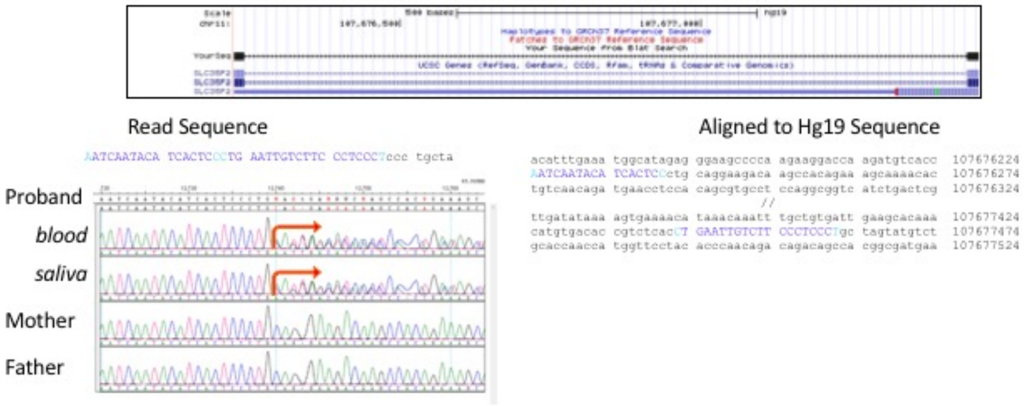


### SERINC5

PCR primers: 2F and 3R
capillary primer: 2F

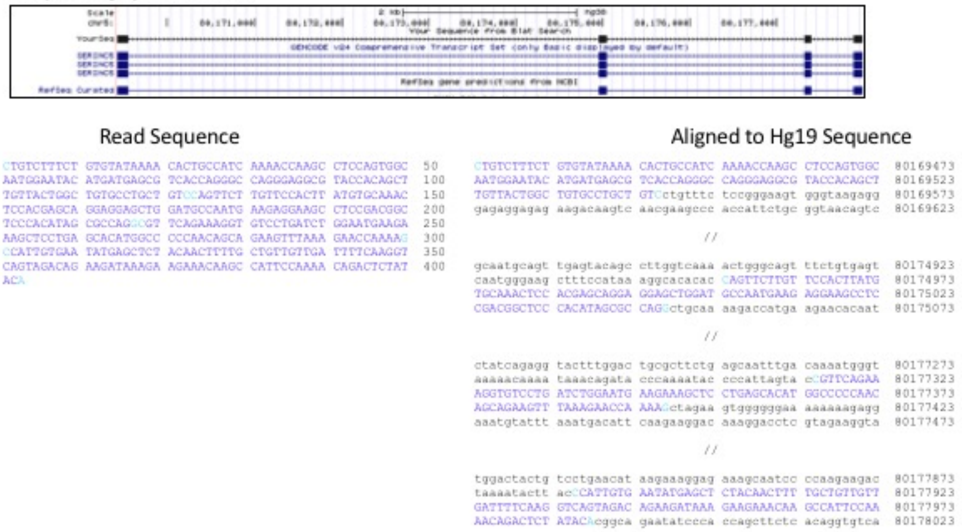
